# Development and retinal remodeling in the brown anole lizard (*Anolis sagrei*)

**DOI:** 10.1101/2021.10.07.462409

**Authors:** Ashley M. Rasys, Shana H. Pau, Katherine E. Irwin, Sherry Luo, Hannah Q. Kim, M. Austin Wahle, Douglas B. Menke, James D. Lauderdale

**Author notes:** Corresponding author and person to whom request should be addressed*: James D. Lauderdale, Department of Cellular Biology, University of Georgia, Athens, GA 30602, USA, Douglas B. Menke, Department of Genetics, University of Georgia, Athens, GA 30602, USA. ***Disclosure Statement***: The authors have nothing to disclose. The funders of this research had no role in study design, data collection and analysis, decision to publish, or preparation of the manuscript.

## Abstract

**Background:** The fovea, a pit in the retina, is believed to be important for high-acuity vision and is a feature found in the eyes of humans and a limited number of vertebrate species that include certain primates, birds, lizards, and fish. At present, model systems currently used for ocular research lack a foveated retina and studies investigating fovea development have largely been limited to histological and molecular studies in primates. As a result, progress towards understanding the mechanisms involved in regulating fovea development in humans is limited and is completely lacking in other, non-primate, vertebrates. To address this knowledge gap, we provide here a detailed histological atlas of retina and fovea development in the bifoveated *Anolis sagrei* lizard, a novel reptile model for fovea research. We also further test the hypothesis that retinal remodeling, which leads to fovea formation and photoreceptor cell packing, is related to asymmetric changes in eye shape.

**Results:** Anole retina development follows the conventional spatiotemporal patterning observed in most vertebrates, where retina neurogenesis begins within the central retina, progresses throughout the temporal retina, and concludes in the nasal retina. One exception to this general rule is that areas that give rise to the fovea undergo retina differentiation prior to the rest of the retina. We find that retina thickness changes dynamically during periods of ocular elongation and retraction. During periods of ocular elongation, the retina thins, while during retraction it becomes thicker. Ganglion cell layer mounding is also observed in the temporal fovea region just prior to pit formation.

**Conclusions:** Anole retina development parallels that of humans, including the onset and progression of retinal neurogenesis followed by changes in ocular shape and retinal remodeling that leads to pit formation in the retina. We propose that anoles are an excellent model system for fovea development research.

**Key Findings:**

- Retina mounding that occurs in foveal areas prior to retinal differentiation progressively disappear as foveal regions of the eye elongate.
- The central and temporal foveal areas undergo retina differentiation before the rest of the retina.
- GCL mounding prior to pit formation occurs in the area of the temporal fovea but not the central fovea.
- When the eye is experiencing ocular retraction, photoreceptor cell packing, and pit formation are observed within foveal regions.

## Introduction

For centuries, physicians and researchers alike have been captivated by the eye and have sought to understand the pathways that enable sight. As a result, decades of work have centered around investigating the development of the human retina and in particular its fovea. Discovered by Samuel Thomas von Sömmerring in the early 1800s, the fovea is a pit located in the macular region of the human eye (Mann, 1928, Barber, 1955) and has long thought to be a critical feature for high visual acuity in humans as well as certain other vertebrates (Slonaker, 1897). Although present in some species of birds (Fite and Rosenfield-Wessels, 1975, Walls, 1942), lizards (Walls, 1942, Underwood, 1970, Röll, 2001, Hulke, 1866), and fish (Collin, 1999, Easter, 1992), much of our understanding of fovea development comes from work in foveated primates (Hendrickson, 1992, Hendrickson, 2015, Hendrickson, 2005, Hendrickson and Drucker, 1992, Hendrickson and Kupfer, 1976, Hendrickson et al., 2012, Hendrickson and Provis, 2006, Hendrickson and Yuodelis, 1984, Springer and Hendrickson, 2005, Springer and Hendrickson, 2004b, Springer and Hendrickson, 2004a, Yuodelis and Hendrickson, 1986, Peng et al., 2019, Voigt et al., 2019, Yan et al., 2020). Histological studies of the primate eye have provided important insights into the timing and progression of retina development and foveal morphogenesis (Hendrickson, 1992, Hendrickson, 2015, Hendrickson, 2005, Hendrickson and Drucker, 1992, Hendrickson and Kupfer, 1976, Hendrickson et al., 2012, Hendrickson and Provis, 2006, Hendrickson and Yuodelis, 1984, Springer and Hendrickson, 2005, Springer and Hendrickson, 2004b, Springer and Hendrickson, 2004a, Yuodelis and Hendrickson, 1986). The fovea develops initially through the lateral displacement of the retina’s ganglion cell layer (GCL) and inner nuclear layer (INL) (Hendrickson and Yuodelis, 1984, Hendrickson et al., 2012). Over time, the photoreceptor cells, which make up the entire outer nuclear layer (ONL), move and pack in around the foveal center (Hendrickson and Yuodelis, 1984, Hendrickson et al., 2012, Yuodelis and Hendrickson, 1986, Hendrickson and Drucker, 1992).

At present, the underlying developmental mechanisms that are involved in reshaping of the retina landscape are not well understood. Nor is it clear how specific retinal regions are instructed to develop a fovea in the first place. Addressing these questions requires the ability to manipulate gene function and/or disrupt cell signaling pathways that have been implicated in fovea development. This is challenging to do in primates because of their reproduction schedule, generation time, and ethical concerns regarding genome modification and the raising of mutant lines. Determining the mechanisms involved in pit formation are also not possible in other animals commonly used for eye research such as the mouse, chick, *Xenopus*, and zebrafish because they all lack a foveated retina. Hence, a new foveated model system is needed--one that can easily be reared in a laboratory setting, reproduces frequently, has a short generation time, and in which the genome can easily be edited. Therefore, we are developing the brown anole lizard (*Anolis sagrei*) as a foveated model system for research (Rasys et al., 2021a, Rasys et al., 2019a, Rasys et al., 2019b, Rasys et al., 2021b). This lizard has a bifoveated retina, possessing a large prominent central fovea and second, much shallower, temporal fovea (Rasys et al., 2021a, Fite and Lister, 1981, Makaretz and Levine, 1980, Sannan et al., 2018, Underwood, 1970, Walls, 1942). Remarkably, during embryonic development anole eyes undergo dynamic changes in their ocular shape (Rasys et al., 2021a). First, the eye dramatically elongates in the regions that will develop a fovea. This is followed by a retraction period, where the eye returns back to its original spherical shape. It is during this retraction phase that the foveae develop. We previously proposed that remodeling of the retina landscape results from changes in eye shape that eventually lead to the organization of high visual acuity areas (Rasys et al., 2021a). Here we test this hypothesis by making a careful examination of the morphology of the retina prior to, during, and after ocular elongation.

## Results

### Structure of the adult retina

The results of this study are made with reference to the retinal structure of the adult anole, which is, as depicted in this sketch made from histological sections cut through anole eyes (Fig. 1), similar to that of other vertebrates. Ganglion cells and putative displaced amacrine cells comprise the ganglion cell layer (GCL). Amacrine cells, bipolar cells, Müller glia, and horizontal cells populate the inner nuclear layer (INL), and photoreceptor cells populate the outer nuclear layer (ONL). The inner plexiform layer is composed of synaptic connections between neurons in the GCL and INL, and is located between these two cell layers. Similarly, the outer plexiform layer is composed of synaptic connections between bipolar cells, horizontal cells and photoreceptors, and is located between the INL and ONL. Immediately adjacent to the ONL on the posterior side of the eye is the retinal pigmented epithelium (RPE). The RPE is a single cell layer with an extensive array of microvilli extending from its apical surface that interdigitates with the outer segments of the photoreceptors. Immediately adjacent to the RPE is the heavily vascularized choroid and cartilaginous sclera, which forms the tough outer surface of the ocular globe.

**Figure 1:**
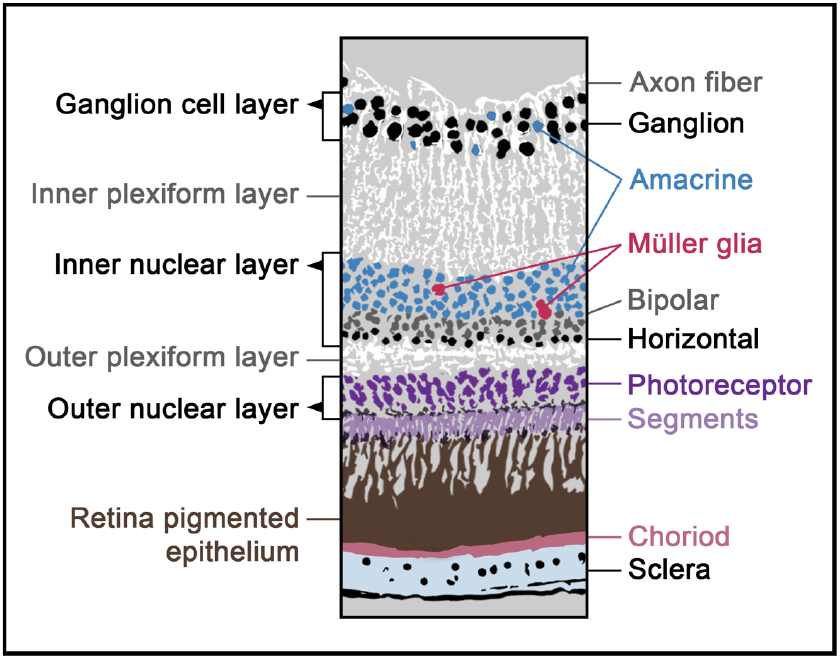
Diagram of cell organization in the adult brown anole lizard retina.

### Early retinal development of foveal areas

In the early developing eye, the prospective central and temporal foveal regions of the neural retina exhibit a thickened and mounded appearance compared to adjacent regions as visualized by histological sections cut through the eyes of stage 5 embryos (Fig. 2a-f; compare regions d,f to b,c,e). These observations suggest that the regions of the retina that will give rise to the central and temporal fovea, respectively, are further along in development than non-foveated retinal regions.

**Figure 2.**
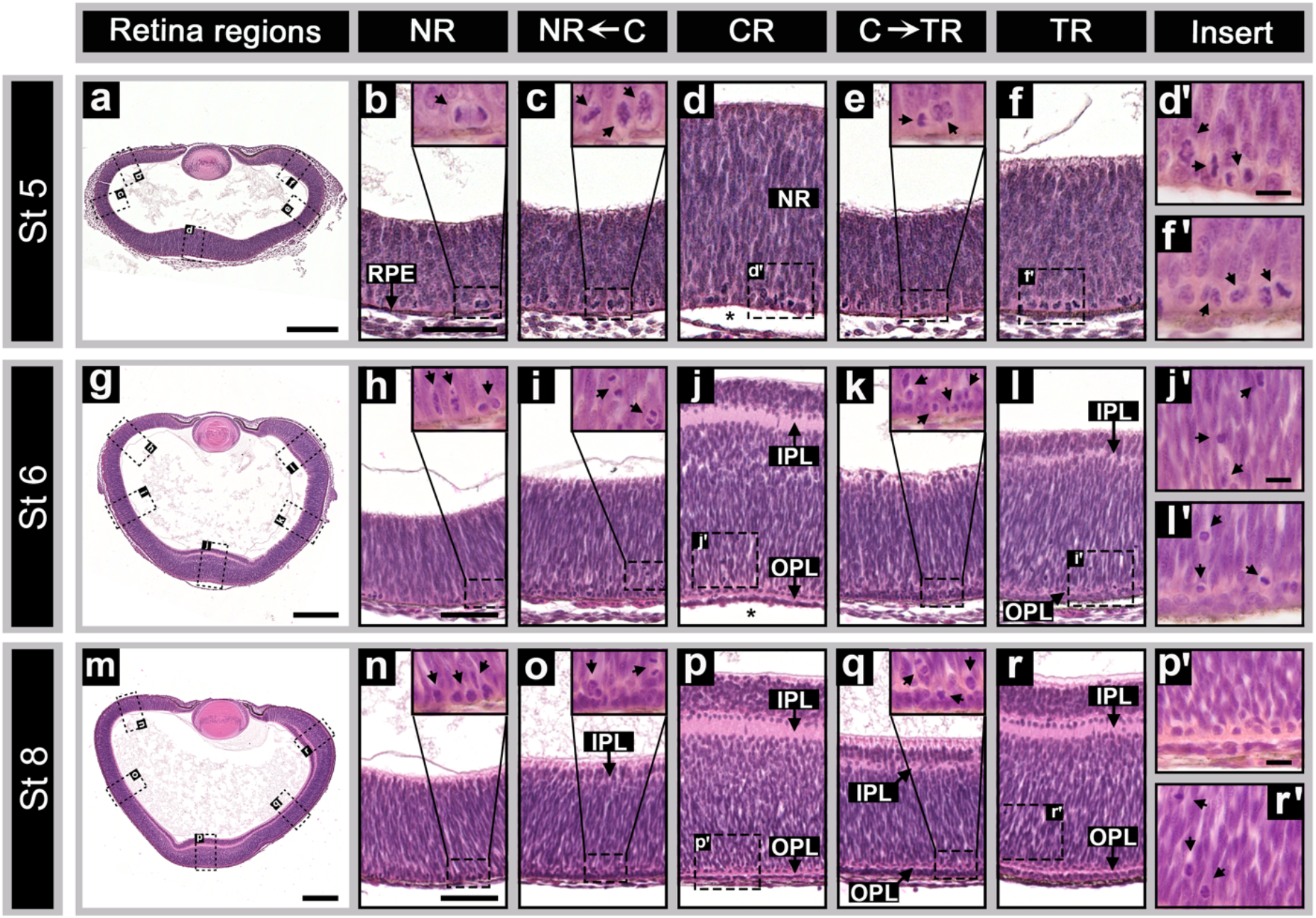
Foveal areas in anoles have retina mounding and undergo retinal lamination before the rest of the retina. Side panels show H&E stained horizontal sections of stage 5 (a), stage 6 (g), and stage 8 (m) embryos. The retina areas: NR—nasal retina (b, h, and n), NR←C—area between the central and nasal retina (c, i, and o), CR—central retina (d, j, and p), C→TR—area between the central and temporal retina (e, k, and q), and TR--temporal (f, l, and r) are shown for each (a), (g), and (m) sections. Magnified inserts of the foveal regions are shown for stage 5 (d’ and f’), stage 6 (j’ and l’), and stage 8 (p’ and r’) embryos. Images (b-f, h-l, and n-r) and (d’-f’, j’-l’, and p’-r’) are respectively to scale with one another. Markers: black arrows – mitotic cells; RPE – retinal pigmented epithelium; NR – neural retina; IPL – inner plexiform layer; OPL – outer plexiform layer; scale bars – 250 μm (a, g, and m), 50 μm (b-f, h-l, and n-r), and 20 μm (d’-f’, j’-l’, and p’-r’).

### Cell proliferation & retina lamination

During the time period that immediately follows optic cup formation in the developing anole (Sanger stages 1-3) (Sanger et al., 2008b) mitotic cells are distributed widely throughout the developing neural retina adjacent to the retinal pigmented epithelium (data not shown; see Fig. 2 of Rasys et al. (2021b)). By late stage 3, an increase in the number of cells dividing in the central region of the retina is observed, and by stage 4, a slight mounding to the retina (*i.e*., increased neural retina thickness) is detected centrally (data not shown; see *Anolis Eye Development* poster from Rasys et al. (2021a)). As the embryo enters stage 5 of development, the eye has grown considerably in overall size and regional differences in both morphology and the relative numbers of mitotic figures become clearly evident (Fig. 2a-f). Retinal mounding in the central retina has increased in both area and thickness and a similar mounded formation, although smaller, is also observed in the temporal region where the presumptive temporal fovea will develop (Fig. 2a, d, f). The highest densities of mitotically active cells are located in the mounded regions of the central and temporal retina (Fig. 2a, d, f), with a lower density of mitotically active cells in the nasal peripheral retina (Fig. 2a-c) and the retinal area between the mounded regions of the central and temporal retina (Fig. 2a, e). Most of the mitotic figures observed in the neural retina are consistent with vertically oriented cell division (the plane of cell division was perpendicular to the plane of the neuroepithelium). In the mounded area of the central retina, cells dividing horizontally are also observed (Fig. 2d).

The first signs of retina lamination are observed by stage 6. A prominent inner plexiform layer (IPL) and developing outer plexiform layer (OPL) are found centrally, while temporally both of these plexiform layers are present but less well developed (Fig. 2g, j, l). Strikingly, no lamination is evident between the central and temporal retina regions (Fig. 2k), nor is lamination observed in the temporal anterior margin. This provides evidence that areas with retinal mounding have progressed further in development as compared to other regions of the retina. During this period, cell proliferation has largely ceased in the central retina and is greatly reduced in the temporal retina. The few cells that are still observed to be undergoing mitosis within these areas are located distally from the ventricle surface (Fig. 2j, j’, l, and l’). In contrast, the densities of mitotic figures located in the peripheral retina and in the region between the central and temporal foveal areas of stage 6 embryos are higher compared to the same regions in stage 5 embryos. For these areas, dividing cells are mostly located adjacent to the ventricle surface (Fig. 2h,I,k), but distally located mitotic cells are observed in portions of the peripheral retina starting to form neuropil (Fig. 2i,k).

By stage 8, the IPL and OPL are now observed in the retina between the central and temporal areas but do not extend to the temporal anterior margin (Fig. 2m-r). At this stage, cell proliferation is largely restricted to the anterior nasal retina (Fig. 2n-o). By stage 10, the IPL and OPL extend from the anterior margin on the temporal side of the eye around the retina, but do not quite reach the anterior margin of the eye on the nasal side (Fig. 3e). Proliferation at this time has largely ceased. By stage 11-12, the IPL and OPL extend all the way around the retina (data not shown, but see Fig. 3i for stage 13).

**Figure 3.**
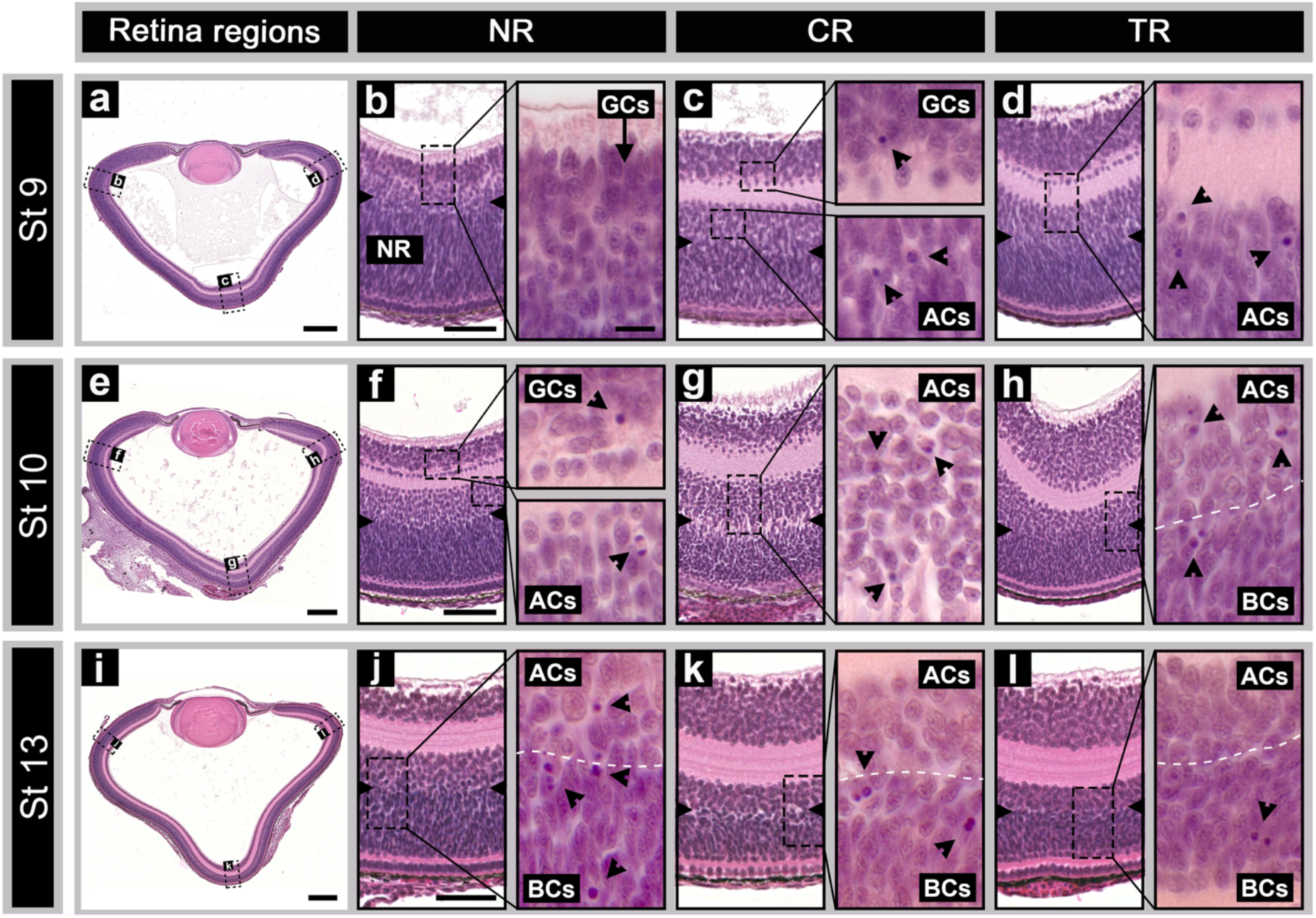
Retinal development in midstage embryos. Panels to the left show horizontal H&E stained sections of stage 9 (a), stage 10 (e), and stage 13 (i) embryos. The nasal retina is depicted in (b, f, and j), the central retina in (c, g, and k), and temporal retina in (d, h, and l). Markers: black arrows – presumptive horizontal cells; black arrow heads – pyknotic cells; side notches – presumptive amacrine/bipolar boundary; NR – neural retina; GCs – ganglion cells; ACs – amacrine cells; BCs – bipolar cells; dashed white lines – presumptive amacrine/bipolar boundary; and scale bars – 250 μm (a, e, and i), 50 μm (b-l), 20 μm (magnified inserts). Inserts are all to scale with each other.

### Neurogenesis of retina cell-types

We next sought to determine the approximate timeline of when cell-types emerge in the developing anole retina. In other vertebrates, the first retina cells to be born are ganglion cells followed by horizontal, amacrine, and cone photoreceptor cells. In contrast, rod photoreceptors, bipolar, and Müller glia cells differentiate later in retina development (Prada et al., 1991, Kahn, 1974, Weysse and Burgess, 1906, Coulombre, 1955, Rhodes, 1979, La Vail et al., 1991, Smith et al., 2001). We monitored the appearance of these cell types in *A. sagrei* following morphological criteria described in other histological studies of the retina (O’Rahilly and Meyer, 1959, Weysse and Burgess, 1906, Coulombre, 1955). As retinal cells differentiate, their cellular morphology changes from spindle (progenitor cell) to round (differentiated) shape (Weysse and Burgess, 1906, O’Rahilly and Meyer, 1959, Coulombre, 1955, Kahn, 1974, Rhodes, 1979). In the anole, the first morphological indication of ganglion cell presence is observed beginning at stage 5 in the central and temporal mounding areas of the retina. Post-mitotically these cells are large, round and are located at the vitreal surface of the neural retina (Fig. 4b, c). Axon fibers extending from these cells toward the future optic stalk are also just becoming apparent. At stage 6, the GCL is clearly visible in the mounded regions of the central and temporal retina (Fig. 2g,j,l; Fig. 4e,f). The GCL is 5-6 cell bodies deep in the central retina and 2-3 cells deep in the temporal retina (Fig. 4e,f). Outside of these two regions, ganglion cells are present but are few in number (*e.g*. Fig. 4d). By stage 9 the GCL is approximately 7-8 cells deep (in the central and temporal retina) and 2-3 cells deep in the retina between these two regions. Nasally, the GCL is just becoming morphologically distinct (Fig. 4g). Shortly after this period, the central and temporal areas of the developing retina become notably longer concordant with asymmetric expansion of the ocular globe along the nasotemporal and lateromedial axes compared to the dorsoventral axis (St 9-14) (Rasys et al., 2021a). This change in length is accompanied by a regional decrease in the cellular density of the neural retina, which is noticeable as a thinning of the GCL (Rasys et al., 2021a). By stage 15 the ocular globe begins to retract, concomitant with regional increases in cellular density (Rasys et al., 2021a). By the time of hatching, the GCL is 4-6 cells deep in all retinal regions except that of the central fovea, which is devoid of ganglion cells due to lateral displacement (Fig. 4m-o).

**Figure 4.**
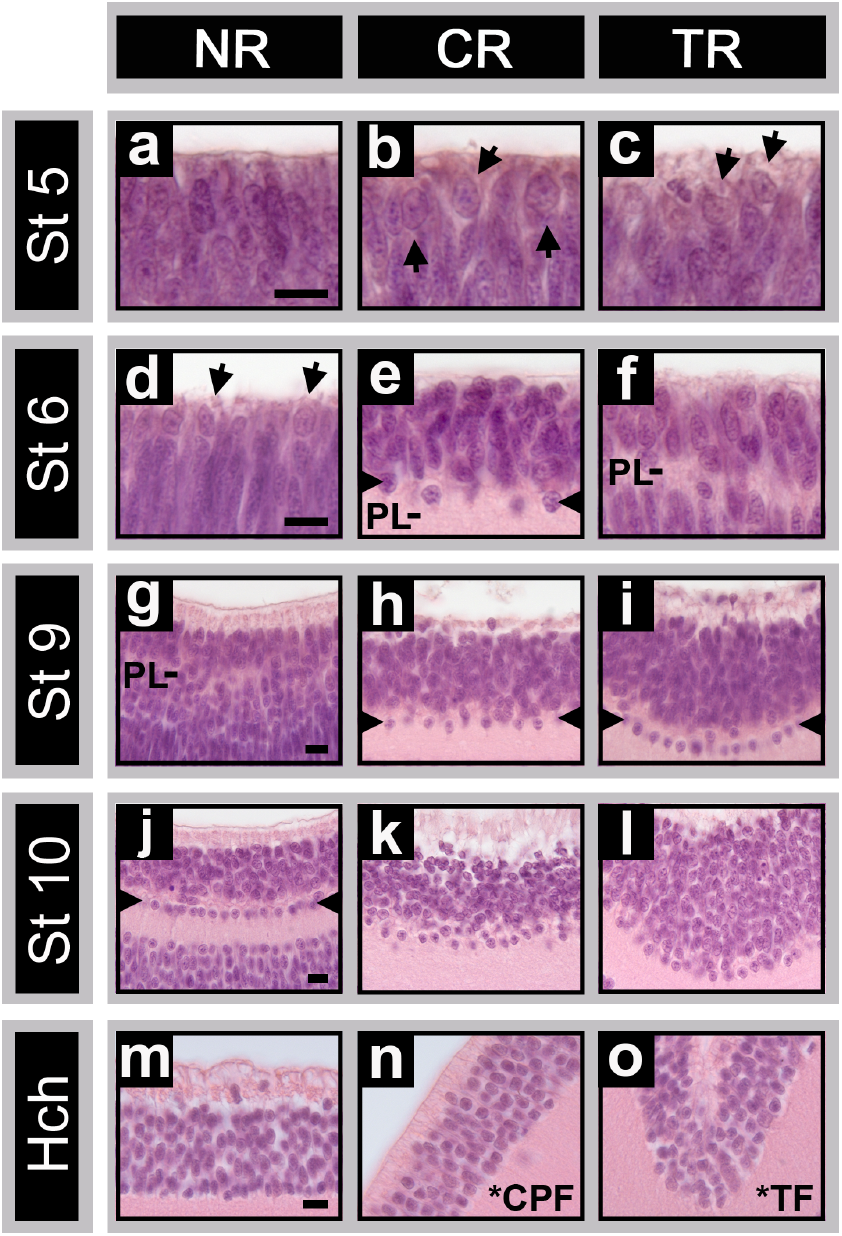
Maturation of the anole retina ganglion cell layer. Horizontal H&E stained sections of stage 5 (a-c), stage 6 (d-f), stage 9 (f-i), and stage 10 (j-l) embryos are shown along with the hatchling (m-o). Images (a, d, g, j, and m) are from the nasal retinal region (NR), (b, e, h, k, and n) from the central region (CR), and (c, f, i, l, and o) from the temporal retina area. Images (a-c), (d-f), (g-i), (j-l), and (m-o) are to scale with each other. Markers indicate: black arrows – presumptive ganglion cells; side notches – organized row of presumptive ganglion or displaced amacrine; PL – plexiform layer; *CPF – central parafoveal area; *TF – temporal fovea area; and scale bars – 20 μm (a-o).

Interestingly, an apparently transient row of cells is distinctly visible in the IPL adjacent to the GCL for a period of time during retinal development. These cells are first seen in the central retina at stage 6 (Fig. 2j,p), next in the temporal retina in the presumptive foveal region (Figs. 2r; 3d; 4i), and finally in the nasal retina (Figs. 3f; 4j). The timing of regional appearance and subsequent disappearance of these cells is linked with the progression of retinal differentiation within the eye. The identity of these cells is not known at this time, but may be ganglion cells and/or displaced amacrine cells.

Amacrine cells are located in the region of the INL located adjacent to the IPL. Morphologically, differentiated amacrine cells are round in shape with a larger soma than retinal progenitor cells, which exhibit a spindle-like form and are densely packed within the neural retina (Fig. 3). Morphologically distinct amacrine cells first appear in small numbers in the central retina at stage 6 (Figs. 2j; 5a-c), in the temporal retina by stage 7 (data not shown), and in the nasal retina by stage 9 (Figs 3a,b; 5d). The region of the INL enriched for amacrine cells can be distinguished from that enriched for bipolar soma by differences in cell density and soma morphologies between amacrine and developing bipolar cells. These regions are easily distinguished from one another throughout embryonic development and into adulthood (for a clear representation see Fig. 3). Interestingly, we also observed that this boundary transiently becomes more distinct and has a marked separation at or around the presumptive fovea areas between stages 10-12. In this separated area, very few cell bodies are present (Figs. 3e,g; 5h).

Similar to the GCL, the amacrine cell density undergoes regional changes during the period the ocular elongation and retraction. Between stages 8-9, both the central and temporal regions have an amacrine cell layer that ranges from 8-9 cells in depth (Fig. 5e-f). At the height of elongation (St 14), this number drops to around 5-6 cells in depth, but during retraction (St 18), the number of cells ranges from 7-9 in depth throughout the retina (Fig. 5j-m, p). During foveae formation in the central and temporal areas (St 16), the amacrine cell layer thins progressively as cell bodies become laterally displaced from the foveal centers. The displacement of amacrine cells away from foveal centers results in an increase in cell density and retinal thickness in regions adjacent to the foveal areas. By hatching, the central fovea is completely devoid of amacrine cells, while the temporal fovea retains a layer of amacrine cells that is 3-4 cells deep (Fig. 5r; Fig. 6).

**Figure 5.**
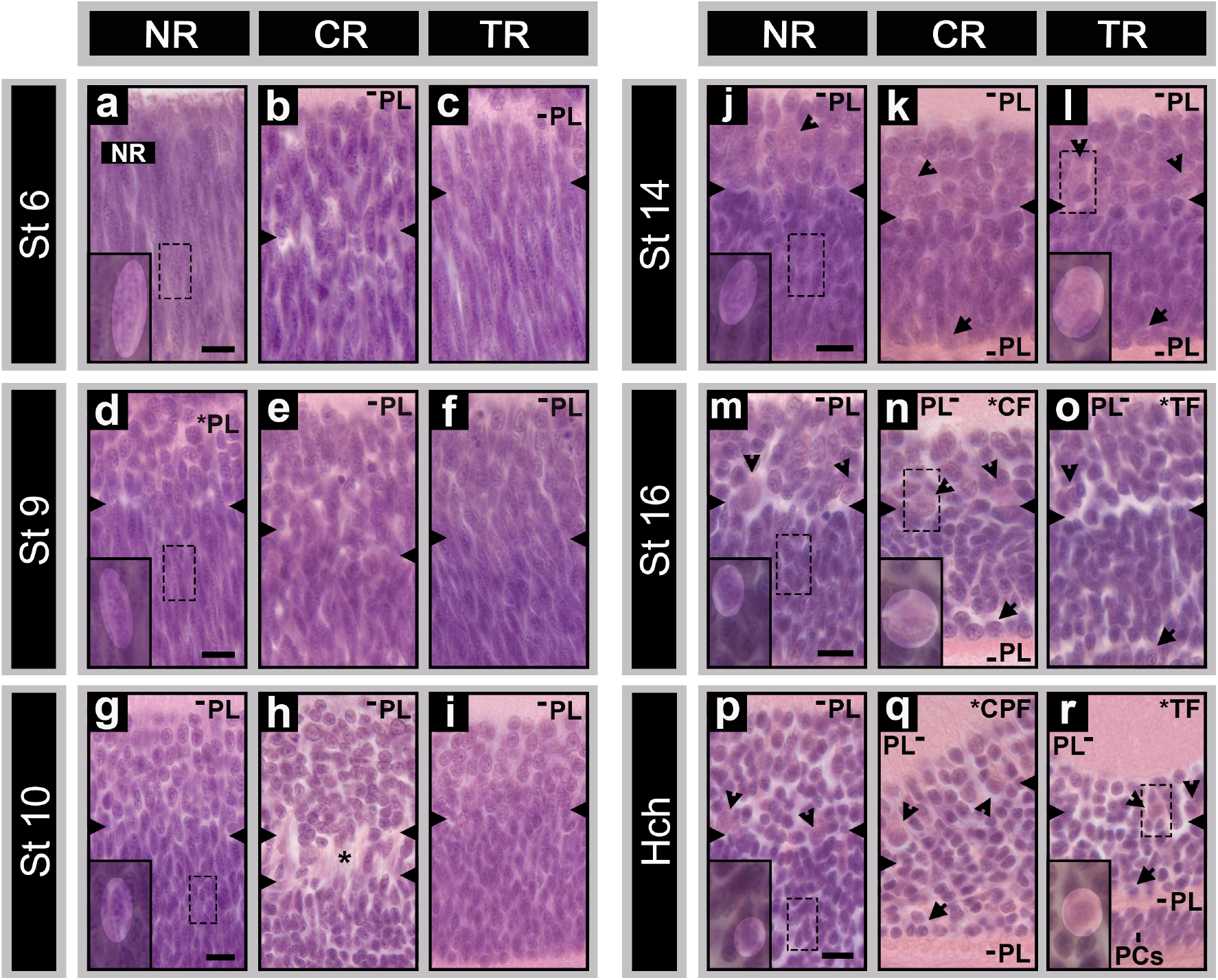
Bipolar and Müller glia histogenesis in the anole retina. Horizontal sections from embryonic stages 6 (a-c), 9 (d-f), 10 (g-i), 14 (j-l), 16 (m-o), and the hatchling (p-r) are shown. Nasal retina regions are shown in (a, d, g, j, m, and p), central retina (b, e, h, k, n, and q), and temporal retina in (c, f, i, l, o, and r). Magnified inserts in NR sections (a, d, g, j, m, and p) show presumptive bipolar cells transitioning from spindle to round morphology. Magnified inserts in CR and TR sections (l, n, and r) depict müller glia cells. Markers: black arrows – presumptive horizontal cells; black arrow heads – presumptive müller glia cells; side notches – presumptive amacrine/bipolar boundary; PL – plexiform layer; asterisk – transient layer of Chievitz; *CF – central fovea; *CPF – central parafoveal region; *TF – temporal fovea; PCs – photoreceptor cells; and scale bars – 20 μm (a-u). Images (a-c), (d-f), (g-i), (j-l), (m-o), and (p-r) are respectively to scale.

**Figure 6.**
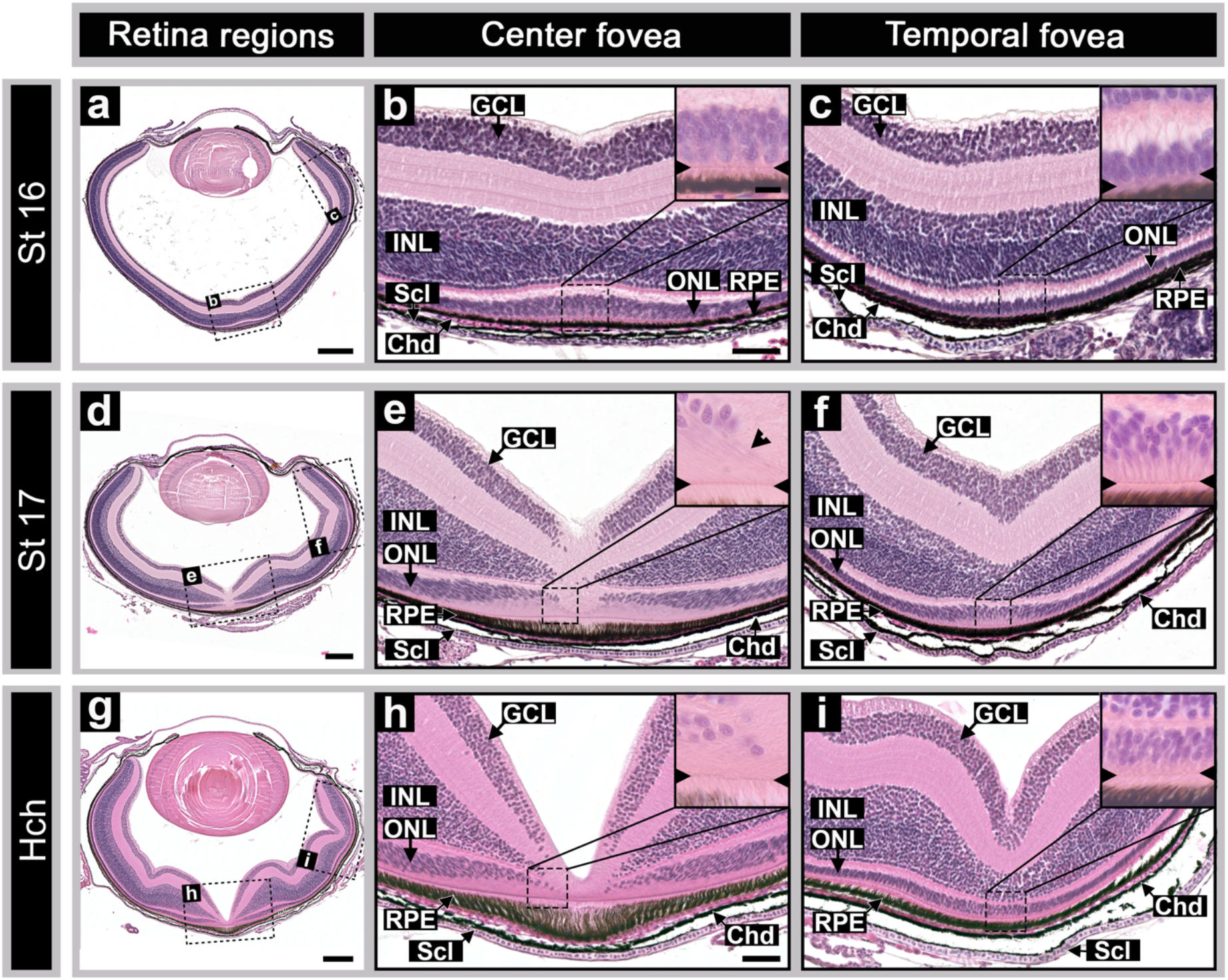
Fovea formation and photoreceptor cell packing in the anole retina. Top panels shows stage 16 embryo (a-c), middle stage 17 embryo (d-f), and hatchling (g-i) horizontal H&E stained sections. Central fovea areas are depicted in (b, e, and h), temporal fovea regions in (c, f, and i). Magnified inserts show photoreceptor cells. Markers signify: GCL – ganglion cell layer; INL – inner nuclear layer; ONL – outer nuclear layer; RPE – retinal pigmented epithelium; Chd – choroid; Scl – sclera; side notches – external limiting membrane; and scale bars – 250 μm (a, d, and g), and 50 μm (b-c, e-f, and h-i). Magnified inserts are all to scale with one another.

Horizonal cells, which occupy the outermost region of the INL, and photoreceptor cells, which lie adjacent to the ventricular surface, are first observed in the central, temporal, and nasal retina areas by stages 5, 6, and 9, respectively (Fig. 7b, e-g). By stage 10, the horizontal cells are neatly arranged into a single row of cells. From this point until hatching, we detect little change either in cellular morphology or number of horizontal cells. The ONL during this time is largely composed of photoreceptors approximately 1-2 soma deep. Interestingly, the distance between neighboring photoreceptor cells is different between the nasal retina and presumptive foveal areas of the central and temporal retina. In the nasal region, cells are closely packed together, while in the central and temporal regions cells are more spaced apart (Fig. 7g-o). This is most evident during the period of ocular elongation (St 9-14). When the eye undergoes retraction (St 15-18), this spacious layout of the central and temporal photoreceptor cells is lost (Fig. 7p-r). By stage 14, the external limiting membrane (ELM) is present along with the first morphological indications of inner photoreceptor cell segments (Fig. 7n and o). By stage 16, the number of photoceptor cells within the central fovea has increased from 1 to 5 cells deep, photoreceptors are 2-3 cells deep in the temporal fovea (Fig. 7q-r). At the time of hatching, photoreceptor and horizontal cells present in the central fovea have become completely laterally displaced, their processes elongated, and inner/outer segments lengthened, whereas in the temporal foveal center photoreceptor cells are 4-5 cell cells deep and horizontal cells are still present (Fig. 7s-u).

**Figure 7.**
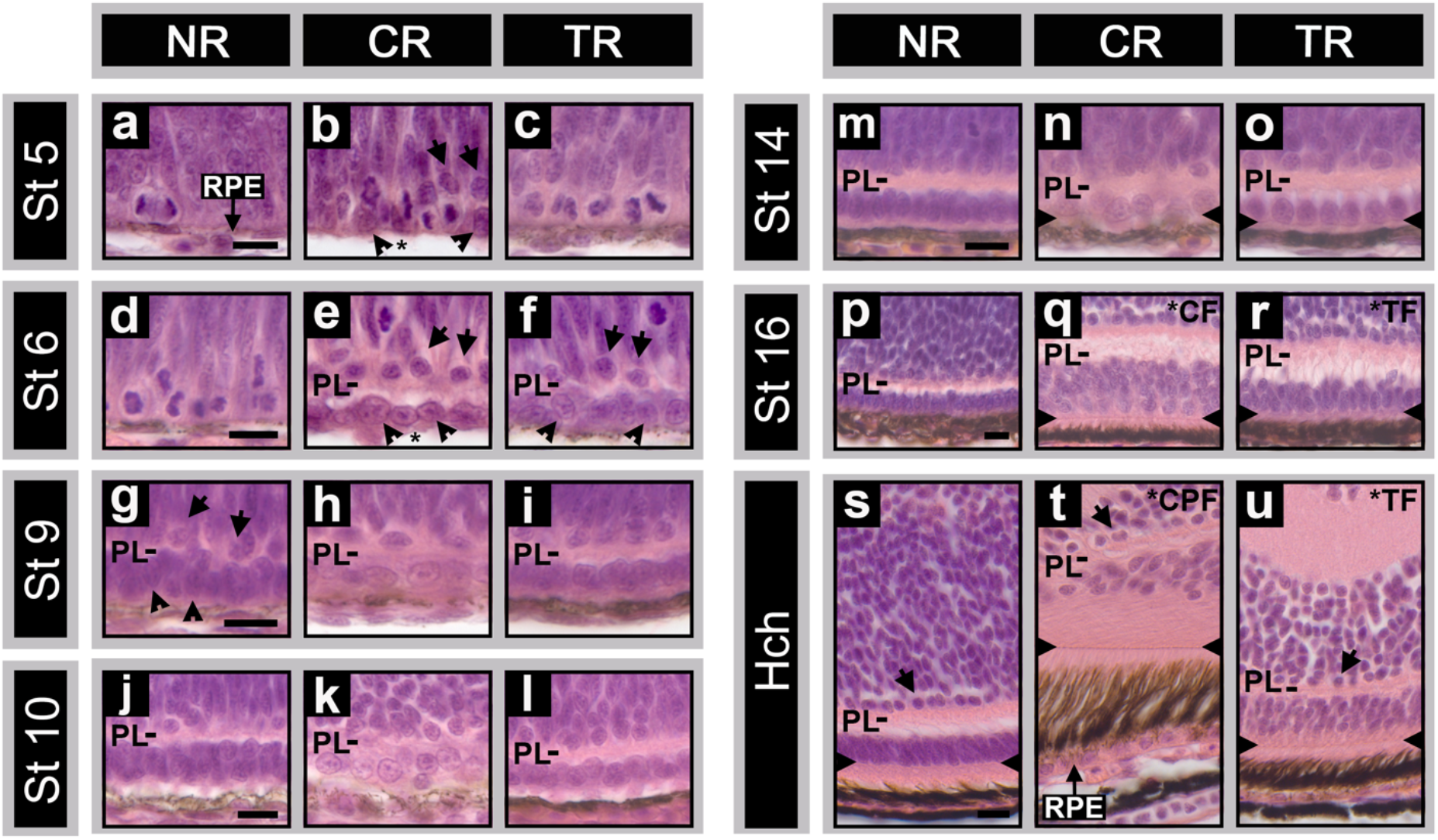
Horizontal and photoreceptor cell development in the anole retina. Images of H&E stained horizontal sections from stage 5 (a-c), stage 6 (d-f), stage 9 (g-i), stage 10 (j-l), stage 14 (m-o), stage 16 (p-r) embryos are depicted along with the hatchling (s-u). Nasal retina regions are shown in (a, d, g, j, m, p, and s), central retina (b, e, h, k, n, q, and t), and temporal retina in (c, f, i, l, o, r, and u). Markers indicate: black arrows – presumptive horizontal cells; black arrow heads – presumptive photoreceptor cells; side notches – external limiting membrane; RPE – retina pigmented epithelium; asterisk – RPE absent due to sectioning artifact; PL – plexiform layer; *CF – central fovea; *CPF – central parafoveal region; *TF – temporal fovea; and scale bars – 20 μm (a-u). Images (a-c), (d-f), (g-i), (j-l), (m-o), (p-r), and (s-u) are to scale with one another, respectively.

Nestled between the amacrine and horizontal cells is a dense population of presumptive bipolar neurons. Due to the lack of definitive morphological features and challenges associated with distinguishing cells in regions of high cell density, we report here only a rough timeline of when observable changes in cell morphology (i.e., a shift from spindle to round morphology) were seen in a large number of cells. Presumptive bipolar cells were first detected between stages 9-10, 10, and 12 for the central, temporal, and nasal retina regions, respectively (Fig. 5d-i). Progression to later stages resulted in the majority of these presumptive bipolar cells transitioning from spindle to round cell morphology throughout the retina and by stage 14, these cells appear largely uniform in shape (Fig. 5j-l). During formation of the central and temporal foveae (St 16), the number of bipolar neurons decrease at the foveal centers in a manner similar to the amacrine cells. By hatching, the central fovea is devoid of bipolar cells whereas the temporal fovea retains bipolar cells (Fig. 5m-r).

Müller glia soma tend to be larger than the cell bodies of other cells in the retina and are typically located in the region of the INL between the amacrine and bipolar soma. Müller glia nuclei are also more eosinophilic than other cells in the INL. Müller glia processes extend the thickness of the retina to both the outer and inner limiting membranes. The presence of cells matching this morphological description are first observed in small numbers between stages 11, 12, and 13 in the central, temporal, and nasal areas and are more easily detected when the ELM appears at stage 14 (Fig. 5l, n, r).

### Cell Death

Naturally occurring waves of cell death that progress in the same order of neuron birth have been observed in other vertebrates (Beazley et al., 1987). In anoles, we found that cell death follows the same spatiotemporal pattern as neurogenesis (i.e., pyknotic cells appear first in the central retina, followed by the temporal retina, then regions immediately flanking the central retina, and lastly the nasal retina (Fig. 3). Cell death is first detected within the GCL and amacrine layer as early as stage 9, becoming more prominent by stage 10 in the central and temporal areas of the retina. Although a pyknotic cells are also present in these cell layers in the nasal retina at this stage, they are relatively few in number (Fig. 3c, d, f-h). By stage 11, the majority of pyknotic cells throughout the central and temporal regions, are found in the amacrine layer, whereas the bulk of dying cells evident in the nasal, regions immediately flanking the central retina are still within the GCL. This progression continues into stages 12-13 where cell death in the central and temporal retina is largely restricted to bipolar neurons. For the other retinal regions, death was predominantly found in the amacrine layer (St 12) and later, the bipolar region (St 13) (Fig. 3j-l). By stage 15, cell death is absent centrally but is still observed throughout the entire temporal retina, limited to the bipolar region. Nasally, only a few of cells in the amacrine layer are detected. By stage 16, cell death is only detected in the nasal retina in bipolar neurons. At stage 17, no cell death is observed. We did not observe any obvious pyknotic cells present in the photoreceptor and horizontal cell layers. Given that these cells represent a small proportion of cells in the retina, it is possible that cell death in these layers went unnoticed because there are fewer cell death events occurring.

### Fovea formation & retinal remodeling

We next turned our attention towards assessing how ocular elongation and retraction contribute to remodeling the retina. We began by looking at the development of the foveae of the retina. Formation of the foveae occur during the last week of embryonic development in anoles, between stages 16-18; it is during this time period that the eye is actively undergoing ocular retraction (Rasys et al., 2021a). As the foveae form, they do so through the lateral displacement of the retina nuclear layers and the movement of photoceptor cells towards the foveal centers. Halfway through stage 16, the first morphological signs of foveae formation can be detected. At the central fovea an indentation in the GCL and a pronounced increase in the number of photoreceptor cells (5 deep) in the ONL is observed (Fig. 6a-c). By stage 17, all three nuclear layers— GCL, INL, and ONL have become laterally displaced at the central fovea. As the displacement of the ONL occurs, the elongated photoreceptor cell processes and the cell bodies appear to extend approximately 45° away from the ELM at the foveal center (Fig. 6e). The lateral displacement of these cells is present well into the parafoveal region during this period. By stage 18 the period ocular retraction is complete, and the central fovea has all of the essential characteristics present in the adult. The temporal fovea develops somewhat differently. Before a shallow pit is observed, the retina briefly increases in width, forming a slight mound or dome (Fig. 6a,c). This doming of the temporal retina appears to primarily be due to the slight increase in number of ganglion cells present with in this region. A slight increase in photoreceptor cells (2-3 deep) is also apparent at the foveal center. By stage 17, photoreceptors are 4-5 cells deep, the INL has narrowed, and a slight pit has formed (Fig. 6f). At the time of hatching, the temporal fovea pit is more pronounced, and the area where photoreceptor cell packing is observed has expanded (Fig. 6g, i).

We next examined the retina more closely for subtle changes that occur before pit formation. Early in development, shortly before on the onset of neurogenesis, we observed that areas where the foveae will develop are thickened and have a mounded appearance. This thickening is detected as early as stage 4 in the central retina (data not shown) and by stage 5 in the temporal retina (Fig. 2a). By stage 6, these areas become more pronounced as the plexiform layers develop (Fig. 2g). It is during this time that ocular elongation begins in the central and temporal regions of the eye (Rasys et al., 2021a). These regions will continue to elongate, peaking around stage 14 (Rasys et al., 2021a). As this process occurs, retinal mounding in the presumptive foveal areas gradually disappears (Figs. 2m; 3a,e,i). By stages 9 and 10, mounding is no longer present (Fig. 3a,e). Nearing the peak elongation (St 13), the entire retina has dramatically thinned (Fig. 3i). After this point, the eye begins to retract; as it does so, the retina thickens, and pit formation occurs.

To further characterize and quantify alterations to the retina during periods of ocular elongation and retraction, we assessed changes in retina length and thickness from embryos during early retina neurogenesis (St 6; n = 4), at the peak of elongation (St14; n = 4), and at hatching (n = 6). Retinas of each sectioned eye were measured from the temporal to nasal ciliary marginal zones to obtain total retina lengths. Because eye size dramatically varies between these three developmental periods, we separated the retina sections from each eye into ten distinct bins to ensure that retina widths, which were measured at the intersection of each bin, were representative of the same region between the different stage embryos. We found that total retina length follows a similar pattern as changes in ocular shape. During early development (St 6), the mean retina length is 3706 ± 410 μm. When ocular elongation is at its peak, retina length increases to 5063 ± 225 μm. By hatching, when the eye is retracted, retina length has decreased down to 3509 ± 90 μm. We next looked at retina thickness distribution across these timepoints (Fig. 8). We determined that retina thickness largely changes as a function of retina length or development of the plexiform layers. Although stage 6 and stage 14 embryos have similar overall retina thickness (Fig. 8b-c), at stage 14 the plexiform layers are developed, whereas at stage 6 the retina is still differentiating (Fig. 8). As a result, some retina growth is expected to occur from retinal lamination in the stage 6 embryo. From stage 14 to hatching, retina thickness dramatically increases through a thickening of both the nuclear and plexiform layers (Fig. 8c).

**Figure 8.**
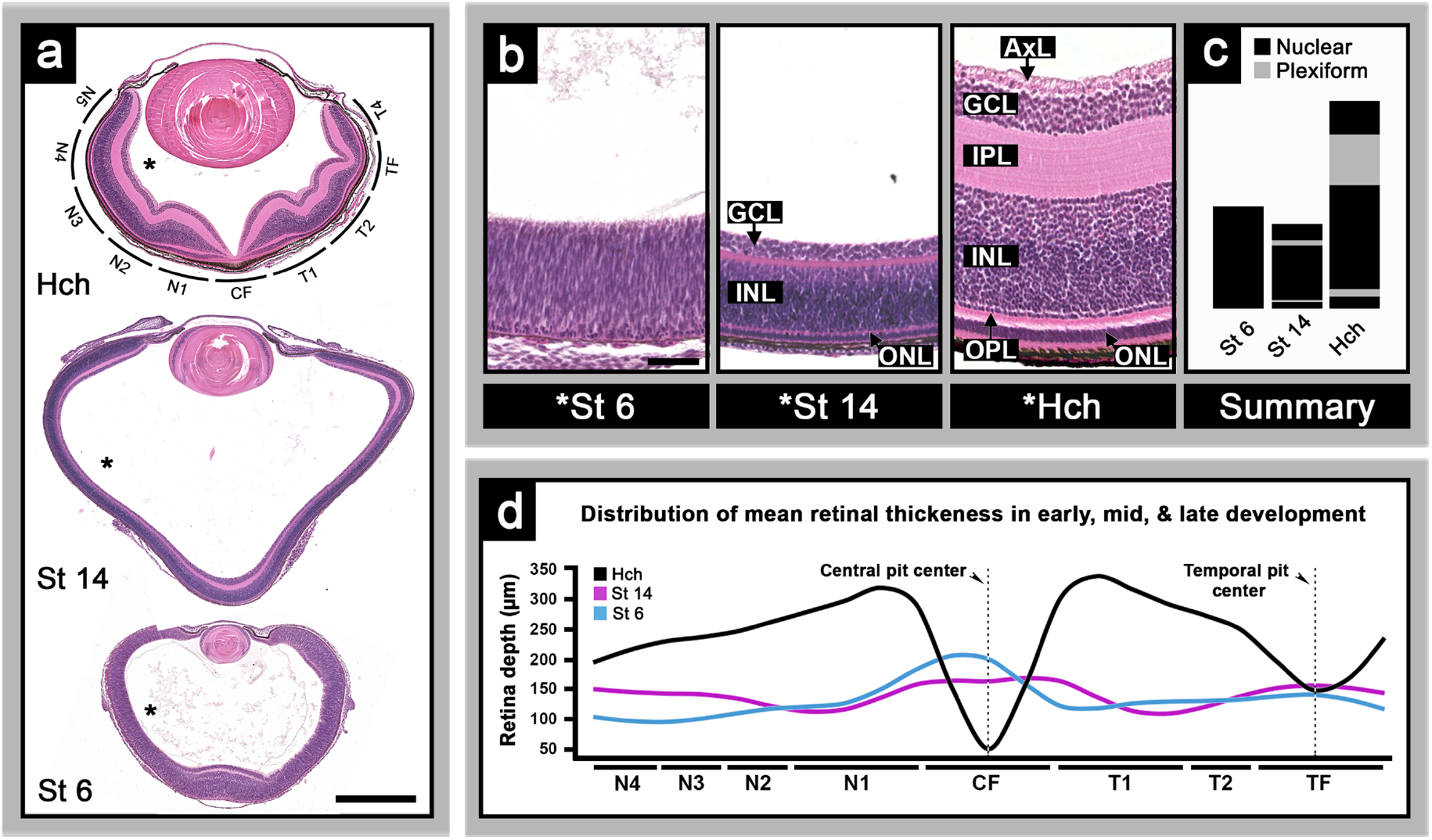
Retinal remodeling during ocular elongation and retraction. Panel (a) shows H&E stained horizontal sections from stage 6 and 14 lizard embryos as well as the hatchling. Retinal binning is depicted around hatchling section shown in (a). Panel (b) shows retina thickness across stages 6, 14 and the hatchling taken from the nasal retina, asterisk area shown in (a). (c) is a diagram summary of (b) comparing nuclear (black) and plexiform (grey) layers. Graph (d) shows mean retina thickness and distribution across embryonic stages 6 (blue), 14 (magenta), and the hatching (black). Markers: GCL – ganglion cell layer; INL – inner nuclear layer; ONL – outer nuclear layer; IPL – inner plexiform layer; OPL – outer plexiform layer; AxL – axon layer; asterisk – retina areas shown in (b); intersection of retina bins (N5, N4, N3, N2, N1, CF, T1, T2, TF, T4) – demarks where retina width measurements were performed; and scale bars – 500 μm (a) and 50 μm (b).

## Discussion

The work we present here shows that foveal regions are morphologically apparent very early on, even before retinal layers are present. We find that differentiation begins first within these regions before the rest of the retina (for illustration see Fig. 9). We also provide evidence that ocular elongation occurs shortly after retina neurogenesis appears complete within these foveal areas. Moreover, consistent with our hypothesis we observed that the retina appears to be influenced by changes in ocular shape and size. As the eye elongates, foveal areas progressively thin and retina mounding is lost. By the point of maximum elongation, when eye size is also at its largest, retina thickness is at its thinnest throughout the entire eye. The eye then retracts, at which time retina thickness increases and remodeling of the retina results in pit formation and photoreceptor cell packing. The timing and sequence of these events—neurogenesis leading to ocular elongation followed by retraction/retinal remodeling (i.e., pit formation and photoreceptor cell packing; for illustration see Fig. 10), suggests that fovea morphogenesis includes a series of steps that begin early and continue throughout most of embryonic development.

**Figure 9.**
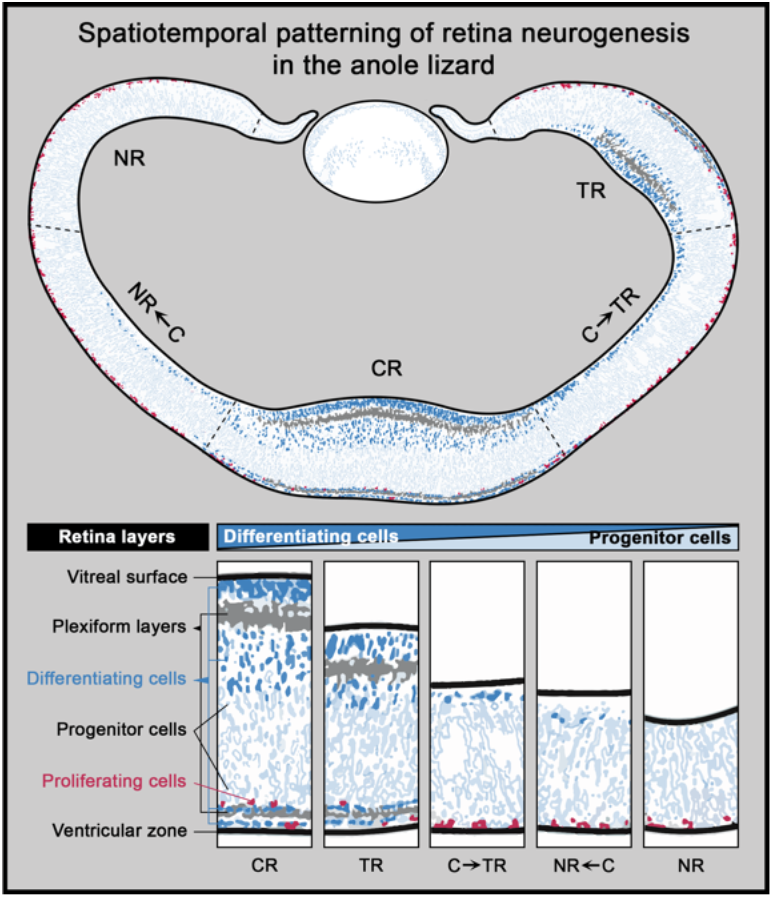
Illustration of the anole’s spatiotemporal patterning of retina neurogenesis. Retina areas: NR – nasal retina; NR←C – retina region between nasal and central retina; CR – central retina; C→TR – region of the retina between central and temporal areas; TR – temporal retina.

**Figure 10.**
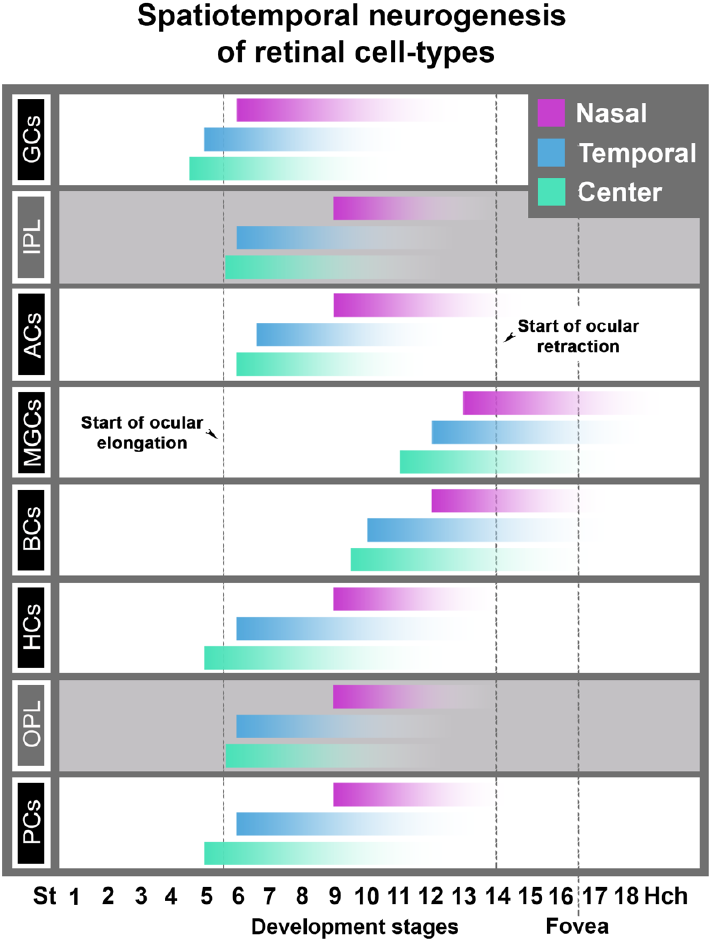
Diagram illustrating the emergence of the different retina cell-types in the anole retina. Nasal (magenta), central (cyan), and temporal (blue) retina patterns are illustrated. Dashed lines indicated periods of ocular elongation, retraction, and pit formation.

Evidence for a relationship between timing of retinal definition and ocular elongation also exists in humans. For instance, during early eye development, the macular region, where the fovea will eventually develop, is also the site where retina neurogenesis first begins and concludes. In humans, this happens between 2-4 months of gestation in the macular region (Rhodes, 1979, Mann, 1928, Hollenberg and Spira, 1973, Barishak, 2001, Hollenberg and Spira, 1972, Hendrickson and Zhang, 2017, Provis et al., 1985, Hendrickson and Provis, 2006, Linberg and Fisher, 1990, van Driel et al., 1990, Hendrickson, 2015, Barber, 1955) and between 5-7 months in the peripheral retina (Barber, 1955, Mann, 1928, Hendrickson, 2015, Hendrickson and Zhang, 2017, Hendrickson, 1992). This 2–4-month time frame is also when asymmetrical changes in ocular shape are observed in human embryonic eyes. For instance, a transient protrusion/bulge develops within the temporal posterior region of the eye where the future fovea develops, starting at around 2 months and peaking at 4 months (von Ammon, 1858, Bach and Seefelder, 1911, Sondermann, 1950, Badtke, 1952, van and Pilleri, 1961). This bulge then progressively disappears between 5-8 months of gestation (von Ammon, 1858, Bach and Seefelder, 1911, Sondermann, 1950, Badtke, 1952, van and Pilleri, 1961, Königstein, 1884), overlapping the time frame when pit formation occurs (Hendrickson and Yuodelis, 1984, Hendrickson et al., 2012, Yuodelis and Hendrickson, 1986, Hendrickson and Drucker, 1992, Mann, 1928). This process in humans is also gradual and first begins with GCL thickening (Hendrickson and Yuodelis, 1984, Hendrickson et al., 2012). This is followed by pit formation, which slowly increases in depth during the latter months of gestation leading up to birth (Hendrickson and Yuodelis, 1984, Hendrickson et al., 2012). Following this step, photoreceptor cells move towards and pack in around the fovea center (Hendrickson and Yuodelis, 1984, Hendrickson et al., 2012, Yuodelis and Hendrickson, 1986, Hendrickson and Drucker, 1992).

Given that the occurrence of asymmetric changes in globe expansion and retraction, one might expect corresponding regional differences in retinal morphology. For instance, when eye elongation occurs, it does so to a much greater extent within the central foveal area compared to the temporal area and the shape of its elongation is also very different (Rasys et al., 2021a). One expected outcome is that we should see differences in retinal remodeling (i.e., pit size and photoreceptor cell packing). Consistent with this, we find that the central pit is more pronounced than that of the temporal fovea and there is a higher degree of photoreceptor packing in the central fovea. Interestingly, the two foveated areas occur through a slightly different sequence of events. In the central fovea, photoreceptor cell packing and retinal lateral displacement occur simultaneously, whereas in the temporal fovea, GCL mounding occurs first, followed by increased number of photoreceptor cells and then pit formation.

In humans, ocular elongation encompasses a broad region of the globe, and like the anole temporal fovea, GCL mounding is observed prior to pit formation (Hendrickson and Yuodelis, 1984, Hendrickson et al., 2012). We believe that GCL mounding before pit formation is an indication of ocular retraction, but the degree of mounding is impacted by the overall shape of ocular elongation. The distribution of forces is different in the central fovea than the temporal fovea. In the central area, elongation is narrow and funnel-like, whereas in the temporal region it is much shallower and broader. If we assume that foveal retina areas are subjected to equal internal resistance (i.e., intraocular pressure) and possess the same intrinsic properties (i.e., equal elasticity), then it is reasonable to expect that mechanical forces exerted at the apex of the elongated regions during retraction would be greater in the central region compared to temporal. For instance, a broader elongated region would likely disperse and alleviate some of the forces generated during ocular retraction whereas a narrower more confined elongated region would tend to focus them. Hence, this could offer some explanation as to why central and temporal foveal areas are morphology different in the anole. Chameleons also have a fovea and undergo similar changes to eye shape (Rasys et al., 2021a). Their central foveal region elongates and retracts to similar degree like anoles, but the shape of its elongation is broader than the anole’s central elongated region. It would be interesting to determine if GCL mounding occurs during central fovea development in chameleons to test this hypothesis.

## Experimental Procedures

### Animals

Lizards are maintained in a laboratory breeding colony at University of Georgia following the anole husbandry and care recommendations outlined by Sanger et al. (2008a). Eggs were maintained in the lab following the protocol previously described by Rasys et al. (2021a). Experiments included the representation of both male and female lizard embryos as well as hatchlings. Hatchlings were euthanized by means consistent with the American Veterinary Medical Association (AVMA) Guidelines for the Euthanasia of Animals.(Association, 2020, Conroy et al., 2009). All experiments were approved, performed, and overseen by the University of Georgia Institutional Animal Care and Use Committee as well as in accordance with the National Institutes of Health Guide for the Care and Use of Laboratory Animals.

### Staging & dissection

Anole embryos develop over a 30-33 day period when incubated at 28°C. Embryos were collected from eggs at various timepoints after egg lay and removed from their egg shells following the protocol described by Rasys et al. (2021a) and staged according to Sanger et al., 2008 guidelines (Sanger et al., 2008b). Eyes were collected from 4 plus embryos from each developmental stage.

### Fixation, embedding, & staining

After dissection, eyes were placed in Bouin’s fixative, which was preferentially chosen over 4% PFA solution because of its excellence at maintaining ocular shape and its resistance to shrinkage artifacts that can occur during the dehydration process. Additionally, it was specifically selected because its quality at preserving individual cell morphology, which enabled easy detection of mitotic, differentiating, and pyknotic cells on histological sections. Eyes were fixed at 4°C overnight on a rocker and then rinsed the next day five times in 1x PBS 15 min washes and then dehydrated in a series of graded ethanol solutions 70%, 80%, 90%, 96% and 100% (twice) for 15 min each. After which, dehydrated eyes were soaked in xylene for 30 min and then incubated in a series of 3 paraffin wax jars for 30 min at 65°C. Eyes were embedded in paraffin, serially sectioned along a horizontal plane at a 10 μm thickness, stained and then mounted following standard hematoxylin & eosin (H&E) and Cytoseal (Thermo Scientific™ Richard-Allan Scientific™) protocols. Sections were imaged using a KEYENCE BZ-700 microscope and photomosaic images were generated with Keyence image stitching software. Contrast and white balance of images were digitally enhanced using Adobe Photoshop CC (2017.01 release).

### Retina measurements

Retina measurements were performed on horizontally sectioned eyes from stages 6 (n= 4), 14 (n= 4), and hatchlings (n= 6) lizards. Total outer and inner retinal lengths were measured and mean plus standard deviation recorded for each section. The outer and inner retina lengths were measured so as to separate the retina into 10 individual bins. Each foveal region was centered in a bin. Retinal width measurements (13 different areas) were performed at the intersection of the bins, foveal center and parafoveal regions. This was done to ensure sampling of the same region across multiple embryo retinas and stages. Retina length and width were recorded in μm units.

## Abbreviations

GCL: Ganglion cell layer
IPL: Inner Plexiform Layer
OPL: Outer Plexiform Layer
INL: Inner nuclear layer
ONL: Outer nuclear layer
ELM: External limiting membrane

## Acknowledgements

The authors wish to thank the members of the Lauderdale and Menke research groups— Rebecca Ball, Christina Sabin, Sergio Minchey, Sukhada Samudra, and Aaron Alcala for all their assistance in the care of Anole lizards at the University of Georgia. The authors also appreciate and thank Drs. Jonathan Eggenschwiler and Heike Kroeger as well as Christina Sabin, and Sukhada Samudra, for their helpful suggestions and editorial comments on this manuscript. The authors also acknowledge the UGA Honors Program and the Center for Undergraduate Research Opportunities, which supported Ms. Katie Irwin, Ms. Sherry Luo, Ms. M. Austin Wahle and Ms. Hannah Kim in the form of CURO Summer Fellowships and CURO Research Assistantships.

## References

Association AVM (2020) AVMA Guidelines for the Euthanasia of Animals: 2020 Edition, American Veterinary Medical Association, Schaumburg, IL.

Bach L, Seefelder R (1911) Atlas zur Entwicklungsgeschichte des menschlichen Auges. Leipzig: W. Engelmann, 1914.

Badtke G (1952) Entwicklungsmechanische Faktoren bei der Formgebung des embryonalen Augapfels. Albrecht von Graefes Archiv für Ophthalmologie, 152, 671–688.

Barber AN (1955) Embryology of the human eye, C. V. Mosby Company, St. Louis, MO.

Barishak YR (2001) Embryology of the Eye and Its Adnexa, Karger.

Beazley LD, Perry VH, Baker B, Darby JE (1987) An investigation into the role of ganglion cells in the regulation of division and death of other retinal cells. Retinal Development, 430, 169–184.

Collin SP (1999) The foveal photoreceptor mosaic in the pipefish, Corythoichthyes paxtoni (Syngnathidae, Teleostei). 14, 369–382.

Conroy CJ, Papenfuss T, Parker J, Hahn NE (2009) Use of tricaine methanesulfonate (MS222) for euthanasia of reptiles. Journal of the American Association for Laboratory Animal Science : JAALAS, 48, 28–32.

Coulombre AJ (1955) Correlations of structural and biochemical changes in the developing retina of the chick. Am J Anat, 96, 153–89.

Easter SS, Jr. (1992) Retinal growth in foveated teleosts: nasotemporal asymmetry keeps the fovea in temporal retina. J Neurosci, 12, 2381–92.

Fite KV, Lister BC (1981) Bifoveal vision in anolis lizards. Brain Behav Evol, 19, 144–54.

Fite KV, Rosenfield-Wessels S (1975) A comparative study of deep avian foveas. Brain, Behav. Evol., 12, 97–115.

Hendrickson A (1992) A morphological comparison of foveal development in man and monkey. Eye (Lond), 6 (Pt 2), 136–44.

Hendrickson A (2005) Organization of the Adult Primate Fovea. In Macular Degeneration (eds Penfold PL, Provis JM), pp. 1–23. Berlin, Heidelberg: Springer Berlin Heidelberg.

Hendrickson A (2015) Development of Retinal Layers in Prenatal Human Retina. 166, 29–35.

Hendrickson A, Drucker D (1992) The development of parafoveal and mid-peripheral human retina. Behav Brain Res, 49, 21–31.

Hendrickson A, Kupfer C (1976) The histogenesis of the fovea in the macaque monkey. Invest Ophthalmol Vis Sci, 15, 746–56.

Hendrickson A, Possin D, Vajzovic L, Toth CA (2012) Histologic development of the human fovea from midgestation to maturity. Am J Ophthalmol, 154, 767–778 e2.

Hendrickson A, Provis J (2006) Comparison of development of the primate fovea centralis with peripheral retina. (eds Harris B, Sernagor E, Wong R, Eglen S), pp. 126–149. Cambridge: Cambridge University Press.

Hendrickson A, Zhang C (2017) Development of cone photoreceptors and their synapses in the human and monkey fovea. J Comp Neurol, 527, 38–51.

Hendrickson AE, Yuodelis C (1984) The morphological development of the human fovea. Ophthalmology, 91, 603–12.

Hollenberg MJ, Spira AW (1972) Early development of the human retina. Can J Ophthalmol, 7, 472–91.

Hollenberg MJ, Spira AW (1973) Human retinal development: Ultrastructure of the outer retina. American Journal of Anatomy, 137, 357–385.

Hulke JW (1866) On the chameleon’s retina; a further contribution to the minute anatomy of the retina of reptiles. Philosophical Transactions of the Royal Society of London,223–229.

Kahn AJ (1974) An autoradiographic analysis of the time of appearance of neurons in the developing chick neural retina. Developmental Biology, 38, 30–40.

Königstein L (1884) Histiologische Notizen. Albrecht von Graefes Archiv für Ophthalmologie, 30, 135–144.

La Vail MM, Rapaport DH, Rakic P (1991) Cytogenesis in the monkey retina. J Comp Neurol, 309, 86–114.

Linberg KA, Fisher SK (1990) A burst of differentiation in the outer posterior retina of the eleven-week human fetus: an ultrastructural study. Vis Neurosci, 5, 43–60.

Makaretz M, Levine RL (1980) A light microscopic study of the bifoveate retina in the lizard Anolis carolinensis: general observations and convergence ratios. Vision Res, 20, 679–86.

Mann IC (1928) The development of the human eye, The University Press.

O’Rahilly R, Meyer DB (1959) The early development of the eye in the chick Gallus domesticus (stages 8 to 25). Acta anatomica, 36, 20–58.

Peng YR, Shekhar K, Yan W, et al. (2019) Molecular Classification and Comparative Taxonomics of Foveal and Peripheral Cells in Primate Retina. Cell, 176, 1222–1237 e22.

Prada C, Puga J, Perez-Mendez L, Lopez R, Ramirez G (1991) Spatial and Temporal Patterns of Neurogenesis in the Chick Retina. Eur J Neurosci, 3, 559–569.

Provis JM, van Driel D, Billson FA, Russell P (1985) Development of the human retina: patterns of cell distribution and redistribution in the ganglion cell layer. J Comp Neurol, 233, 429–51.

Rasys AM, Divers SJ, Lauderdale JD, Menke DB (2019a) A systematic study of injectable anesthetic agents in the brown anole lizard (Anolis sagrei). Laboratory Animals.

Rasys AM, Park S, Ball RE, Alcala AJ, Lauderdale JD, Menke DB (2019b) CRISPR-Cas9 Gene Editing in Lizards through Microinjection of Unfertilized Oocytes. Cell Reports, 28, 2288-+.

Rasys AM, Pau SH, Irwin KE, et al. (2021a) Ocular elongation and retraction in foveated reptiles. Dev Dyn.

Rasys AM, Pau SH, Irwin KE, Luo S, Menke DB, Lauderdale JD (2021b) Anterior eye development in the brown anole, *Anolis sagrei*. bioRxiv, **bioRxiv** doi: 10.1101/2021.02.15.429783, 1–21.

Rhodes RH (1979) A light microscopic study of the developing human neural retina. Am J Anat, 154, 195–209.

Röll B (2001) Gecko vision—retinal organization, foveae and implications for binocular vision. Vision Research, 41, 2043–2056.

Sanger TJ, Hime PM, Johnson MA, Diani J, Losos JB (2008a) Laboratory protocols for husbandry and embryo collection of Anolis lizards. Herpetological Review, 39, 58–63.

Sanger TJ, Losos JB, Gibson-Brown JJ (2008b) A developmental staging series for the lizard genus Anolis: a new system for the integration of evolution, development, and ecology. J Morphol, 269, 129–37.

Sannan NS, Shan X, Gregory-Evans K, Kusumi K, Gregory-Evans CY (2018) Anolis carolinensis as a model to understand the molecular and cellular basis of foveal development. Exp Eye Res, 173, 138–147.

Slonaker JR (1897) A comparative study of the area of acute vision in vertebrates, Ginn & company, Boston.

Smith RS, John SWM, Nishina PM, Sundberg JP (2001) Systematic Evaluation of the Mouse Eye, CRC Press.

Sondermann R (1950) Die Bedeutung der Vererbung für die Entwicklung der Myopie. Albrecht von Graefes Archiv für klinische und experimentelle Ophthalmologie, 151, 200–208.

Springer AD, Hendrickson AE (2004a) Development of the primate area of high acuity. 1. Use of finite element analysis models to identify mechanical variables affecting pit formation. Vis Neurosci, 21, 53–62.

Springer AD, Hendrickson AE (2004b) Development of the primate area of high acuity. 2. Quantitative morphological changes associated with retinal and pars plana growth. Vis Neurosci, 21, 775–90.

Springer AD, Hendrickson AE (2005) Development of the primate area of high acuity, 3: temporal relationships between pit formation, retinal elongation and cone packing. Vis Neurosci, 22, 171–85.

Underwood G (1970) The Eye. In Biology of the Reptilia, Volume 2. Morphology B (eds Gans C, Parsons TS), pp. 1–93. London and New York: Academic Press, Inc.

van Driel D, Provis JM, Billson FA (1990) Early differentiation of ganglion, amacrine, bipolar, and Muller cells in the developing fovea of human retina. Journal of Comparative Neurology, 291, 203–219.

van L, Pilleri G (1961) [Morphological research on the “scleral protuberance”]. Albrecht Von Graefes Arch Ophthalmol, 163, 1–9.

Voigt AP, Whitmore SS, Flamme-Wiese MJ, et al. (2019) Molecular characterization of foveal versus peripheral human retina by single-cell RNA sequencing. Exp Eye Res, 184, 234–242.

von Ammon FA (1858) Die Entwicklungsgeschichte des menschlichen Auges, Peters.

Walls GL (1942) The Vertebrate Eye and Its Adaptive Radiation, The Cranbrook Institute of Science, Bloomfield Hills, Michigan.

Weysse AW, Burgess WS (1906) Histogenesis of the retina. The American Naturalist, 40, 611–637.

Yan W, Peng YR, van Zyl T, et al. (2020) Cell Atlas of The Human Fovea and Peripheral Retina. Sci Rep, 10, 9802.

Yuodelis C, Hendrickson A (1986) A qualitative and quantitative analysis of the human fovea during development. Vision Res, 26, 847–55.

